# SRF-CLICAL: an approach for patient risk stratification using random forest models

**DOI:** 10.1101/2021.06.22.448514

**Authors:** Felipe A. Simao, Reija Hieta, Jane A. Pulman, Gabriele Madonna, Paolo Antonio Ascierto, Rolf Lewensohn, Giuseppe V. Masucci

**Affiliations:** Genevia technologies OY, Tampere, Finland; Melanoma, Cancer Immunotherapy and Development Therapeutics Unit, Istituto Nazionale Tumori IRCCS Fondazione “G. Pascale”, Napoli, Italy; Tema Cancer, Karolinska Hospital and Department of Oncology-Pathology, Karolinska Institutet, Stockholm, Sweden

## Abstract

An important part of good clinical care is identifying which patients have a high likelihood of experiencing adverse outcomes. Similarly, due to the significant impact cancer treatment can have on a patient’s quality of life, it is also important to properly identify which patients are likely to benefit from more aggressive treatment options. As such, models for predictive risk stratification can be extremely useful in clinical decision making. In this paper, we present, Survival Random Forest-Clinical Categorization Algorithm (SRF-CLICAL), a new method for patient risk stratification using random forests for survival, regression and classification. As a proof of concept, we demonstrate this method on two different cohorts of cancer patients.

## 1. Introduction

Prediction of risk at an individual level has been a perennial problem in medical care. Accurate methods for risk prediction can help clinical decision making about disease progression, trajectory and what is the optimal treatment for a given patient and disease combination [1, 2]. Rapid advances in biomedical knowledge and information technology can be leveraged to significantly improve risk predictions in clinical settings. Over recent years, machine learning methods have been successfully employed in a wide range of settings involving highly complex, heterogeneous data sources. Despite these technological advances, most of the currently used methods for clinical risk prediction are still based on simpler regression approaches which can only handle a small number of predictive variables [3]. The aim of this study was to develop an easy to use, flexible, clinical risk prediction method that can be applied to any disease for which significant amounts of clinical, biomarker and/or genomic data is available. SRF-CLICAL builds upon and improves the performance of the original empirical Clinical Categorization Algorithm (CLICAL) [4, 5], while also making the methodology more generally applicable. As a proof of concept, we demonstrate the suitability of RFS-CLICAL to produce a clinical risk prediction model for metastatic melanoma based on clinical data from a cohort of 578 patients for which 8 clinical variables were available. Further validation was done using an external cohort of 118 metastatic melanoma patients [6], and the generalisability of the approach was demonstrated on a cohort of 580 colorectal cancer patients.

## 2. Methods

The selection of clinical variables was performed using univariate Cox proportional hazards (Cox PH) regression models fitted for all available clinical variables with survival time as the outcome variable, as implemented in the R package survival, v. 3.2-3 [7]. Effron approximation was used for handling tied death times. The p-values and hazard ratios of the models were inspected to compare the predictive abilities of the independent variables. A survival random forest (SRF) model was then computed for the data using a selection of clinical variables. The R package randomForestSRC, v. 2.9.3 [8] was used for computing an initial model using a training dataset (80% of the cohort). An optimised SRF model was generated by tuning the parameters mtry and node size for 50, 100, 200, 500 and 1000 trees, using the tune.rfsrc function of the randomForestSRC package, and a starting mtry value of 2. Out-of-bag (OOB) errors of the models are compared and the number of trees with the smallest OOB error (ntree = 1000) was chosen as the ntree value for the optimised SRF model, with optimal mtry = 2 and nodesize = 10 values for the given number of trees being used for producing the final model. The function predictSurvProb of the R package pec, v. 2019.11.03 was used for making survival probability predictions at the 60 month time-point [9]. Risk categories were then defined based on the distribution of 60 month survival probability predictions. The riskRegression package, v. 2020.02.05 was used for plotting time-dependent Receiver Operating Characteristic (ROC) curves [10]. Lastly, Kaplan-Meier plots were generated using the R packages survival, v. 3.2-3 and survminer, v. 0.4.8 with all patients divided into three risk groups based on the SRF-predicted survival probabilities [4, 8]. Ten-fold cross-validation was performed, using the R package caret to produce balanced validation groups [12]. Detailed methods are described in the supplementary materials.

## 3. Results

The time-dependent ROC curves produced for the optimised SRF-model for metastatic melanoma are shown in Figure 1. The curves show very similar AUC values at the 12 (81.2), 24 (79.8) and 36 (81.5) month timepoints, with a slight decrease in performance at the 60 month time point (71.2). These observations are consistent with the high overall mortality of metastatic melanoma. Survival probability predictions and assigned risk categories are shown in Figure 2. The prediction results indicate that the SRF-CLICAL is capable of stratifying metastatic melanoma patients according to predicted risk, and that risk predictions are consistent with the patient mortality observed for this cohort (p < 0.0001).

**Figure 1.**
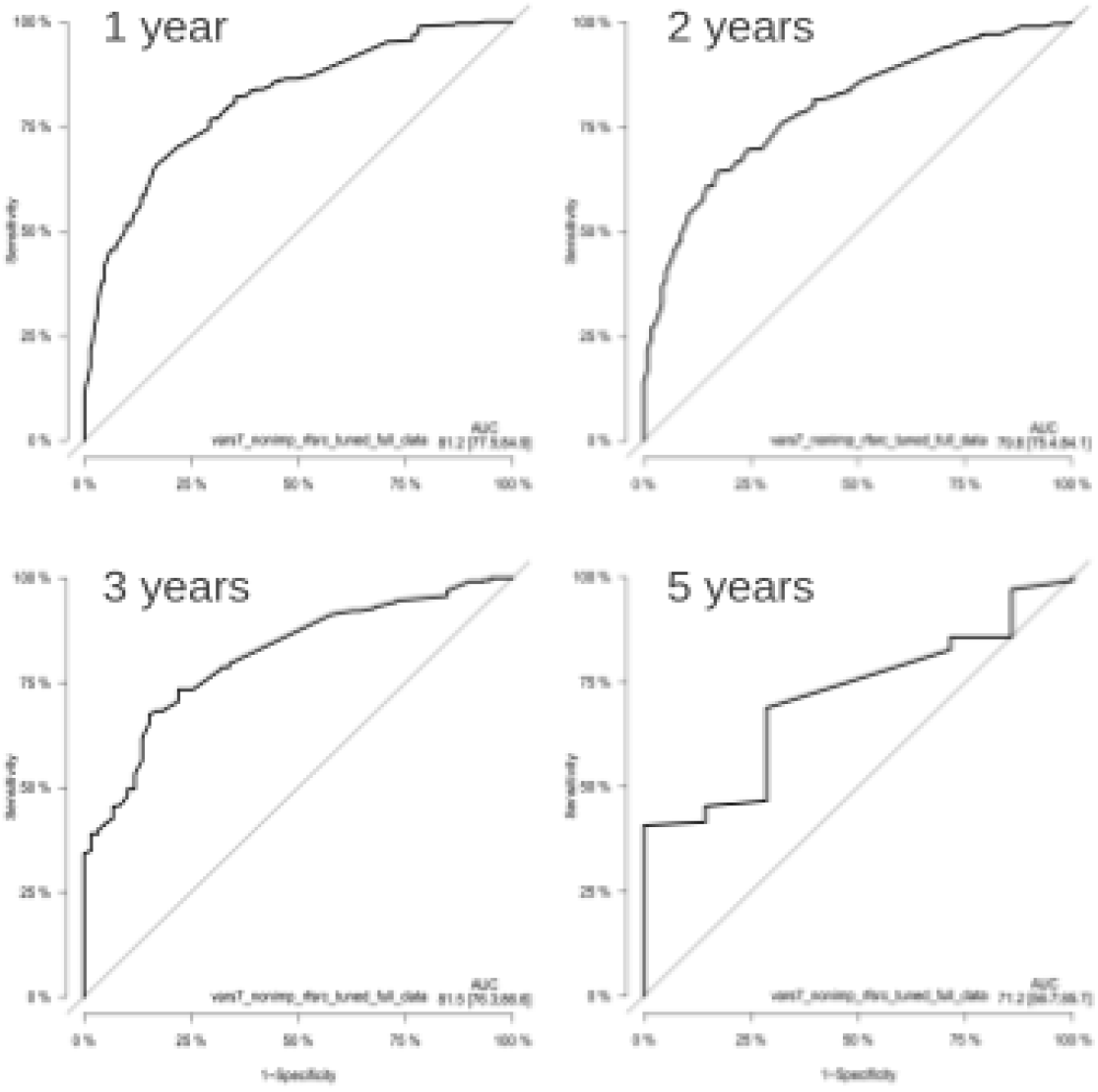
Time-dependent receiver operating characteristic (ROC) curves ot the tuned 7-variable SRF model of the melanoma cohort. ROC curves are shown at time points 12, 24, 36 and 60 months. Y-axis in the plots is displaying the True Positive rate (TPR), i.e. Sensitivity, whereas X-axis in the plots is displaying (the False Positive Rate (FPR), ie. 1-Specificity.

**Figure 2.**
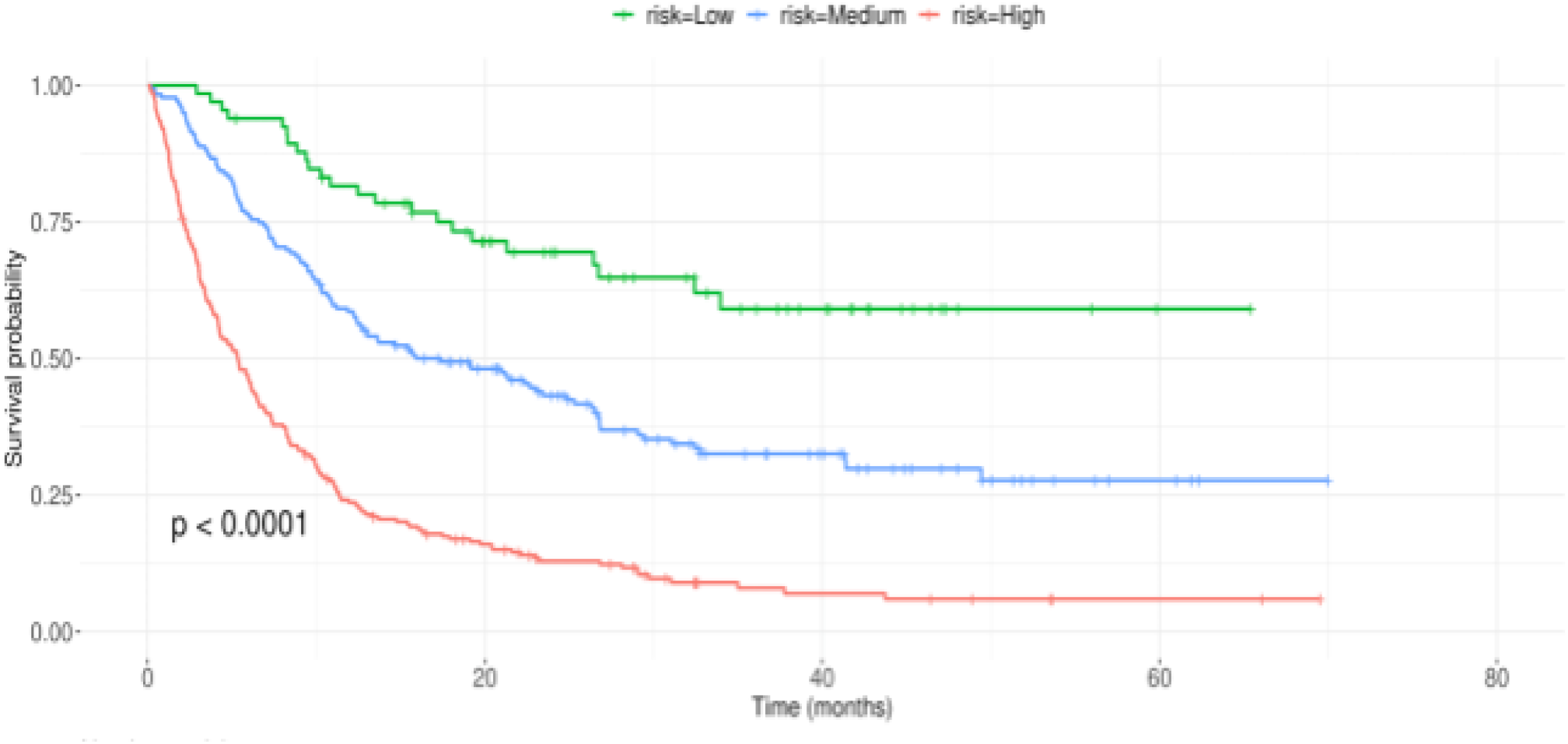
Kaplan-Meier survival curves produced using the survival probability predictions of all 578 melanoma patients divided into three risk groups based on the survival probability predictions. The X-axis shows the survival time in months and Y-axis shows the survival probability estimated based cn the observed and censored events.

## 4. Discussion

The SRF-CLICAL is a new method for clinical risk prediction that builds upon the original empirical CLICAL and existing approaches for survival analysis. SRF-CLICAL uses the R-package Random Forests for Survival, Regression and Classification, as a base for predicting survival probabilities for patients and converts these probabilities into easy to understand individualized clinical risk categories. As demonstrated in our proof-of-concept analysis of the 578 patient metastatic melanoma cohort, the 117 external validation cohort and 580 patient colorectal cancer cohort (Supplementary materials), SRF-CLICAL is particularly suitable for clinical risk prediction in cancer cohorts. Data from any new patients can be used with optimised SRF-models, to predict survival probabilities and assign a risk category. The methodology underlying SRF-CLICAL is generalizable and can be employed with a variety of datasets, regardless of disease type or the number of available variables.

