## Supplementary Materials for "SRF-CLICAL: an approach for patient risk stratification using random forest models"

### 4. Supplementary materials

#### 4.1 Expanded Methods

All eight clinical variables listed in Supplementary Table 1 were subjected to univariate Cox PH analysis. The results of the analysis were combined into a table and ordered based on p-values obtained from the individual analyses (Supplementary Table 2). The results indicated that six of the eight clinical variables under investigation were statistically significantly associated with survival ($p$-value $< 0.05$; blue background), with age group having borderline significance (grey background) and sex being insignificant. To study the effect of the eight different covariates while adjusting for the impact of the others, a multivariate Cox PH model was created. Forest plot of the obtained result is shown in Supplementary Figure 1.

Supplementary Table 1. Clinical variables used in the study

| Variable no | Variable | Groups |
|---|---|---|
| 1 | Sex | Female, Male |
| 2 | Age | Young (<= 60) , Old (> 60) OR Young (<= 65) , Old (> 65) |
| 3 | BRAF | Wild type, Mutated |
| 4 | LDH | Normal, High, Very high |
| 5 | CNS metastasis | No, Yes |
| 6 | Previous treatment | Non-target, Target |
| 7 | Eosinophils | Normal, Elevated |
| 8 | Neutrophils | Normal, Abnormal |

Supplementary Table 2. Results of univariate Cox proportional hazards analyses

| Variable | Beta coefficient | Hazard ratio | 95% conf. interval | Inverse hazard ratio | P-value |
|---|---|---|---|---|---|
| LDH: very high vs normal | 1,395 | 4,035 | 3.1 - 5.3 | 0,248 | 1,75E-24 |
| Neutrophils: abnormal vs normal | 0,680 | 1,974 | 1.6 - 2.4 | 0,506 | 9,72E-12 |
| CNS metastasis: YES vs NO | 0,626 | 1,869 | 1.5 - 2.3 | 0,535 | 1,83E-09 |
| Previous treatment: TARGET vs NON TARGET | 0,510 | 1,665 | 1.4 - 2 | 0,601 | 1,18E-06 |
| LDH: high vs normal | 0,580 | 1,787 | 1.4 - 2.3 | 0,560 | 4,65E-06 |
| Eosinophils: elevated vs normal | -0,438 | 0,645 | 0.44 - 0.94 | 1,549 | 2,32E-02 |
| BRAF: MUT vs WT | 0,224 | 1,251 | 1 - 1.5 | 0,799 | 2,68E-02 |
| Age group: OLD60+ vs young | -0,184 | 0,832 | 0.69 - 1 | 1,202 | 5,92E-02 |
| Sex: M vs F | 0,024 | 1,024 | 0.85 - 1.2 | 0,976 | 8,06E-01 |

A survival random forest (SRF) model was trained using the randomForestSRC, v. 2.9.3 and with seven clinical variables (Age, BRAF, LDH, presence of CNS metastasis, Previous Treatment, Level of Eosinophils and Level of Neutrophils), with sex excluded [8]. In sequence, an optimised SRF model was generated by tuning mtry and node size parameters for 50, 100, 200, 500 and 1000 trees using tune.rfsrc function of the randomForestSRC package, with starting value of mtry set to 2. Out-of-bag (OOB) errors of the models are compared and the number of trees with the smallest OOB error (ntree = 1000) was chosen as the ntree value for the optimised SRF model, with optimal mtry = 2 and nodesize =10 values for the given number of trees used for producing the final model. The R package pec, v. 2019.11.03 [9] function predictSurvProb was then used for making 60 month survival probability predictions for all patients. Time-dependent receiver operating characteristic (ROC) curves at time points 12, 24, 36 and 60 months were then generated (Figure 1). The Area Under the Curve (AUC) values at the first three time points were very similar to each other, around 80, whereas at the 60 month time point the AUC was somewhat lower, 71.2 (Figure 1, bottom right corner).

Kaplan-Meier plots were then generated for patients divided into three risk groups based on the SRF- predicted survival probabilities, using the R packages survival, v. 3.2-3 [7] and survminer, v. 0.4.8 [11]. The full dataset was used for this analysis so that an adequate number of patients could be assigned for each group. Distribution of the predicted survival probabilities at 5 years (60 months) were examined and used for defining the three possible risk categories: patients with survival probability < 0.2 were categorised into as High risk, patients with survival probability => 0.41 were categorised into the Low risk and patients with survival probability in between these thresholds were categorised into the Medium risk. The SRF-predicted survival probabilities were then used to assign one risk category to each of the 578 patients of this cohort. Kaplan-Meier survival curves for these three patient groups were then compared (Figure 2). The three risk groups showed clearly and statistically significantly distinct survival curve profiles, with patients categorised into the High risk group having much lower median survival time (~5 months) than the patient in the Medium (~18 months) and Low risk (> 65 months) groups.

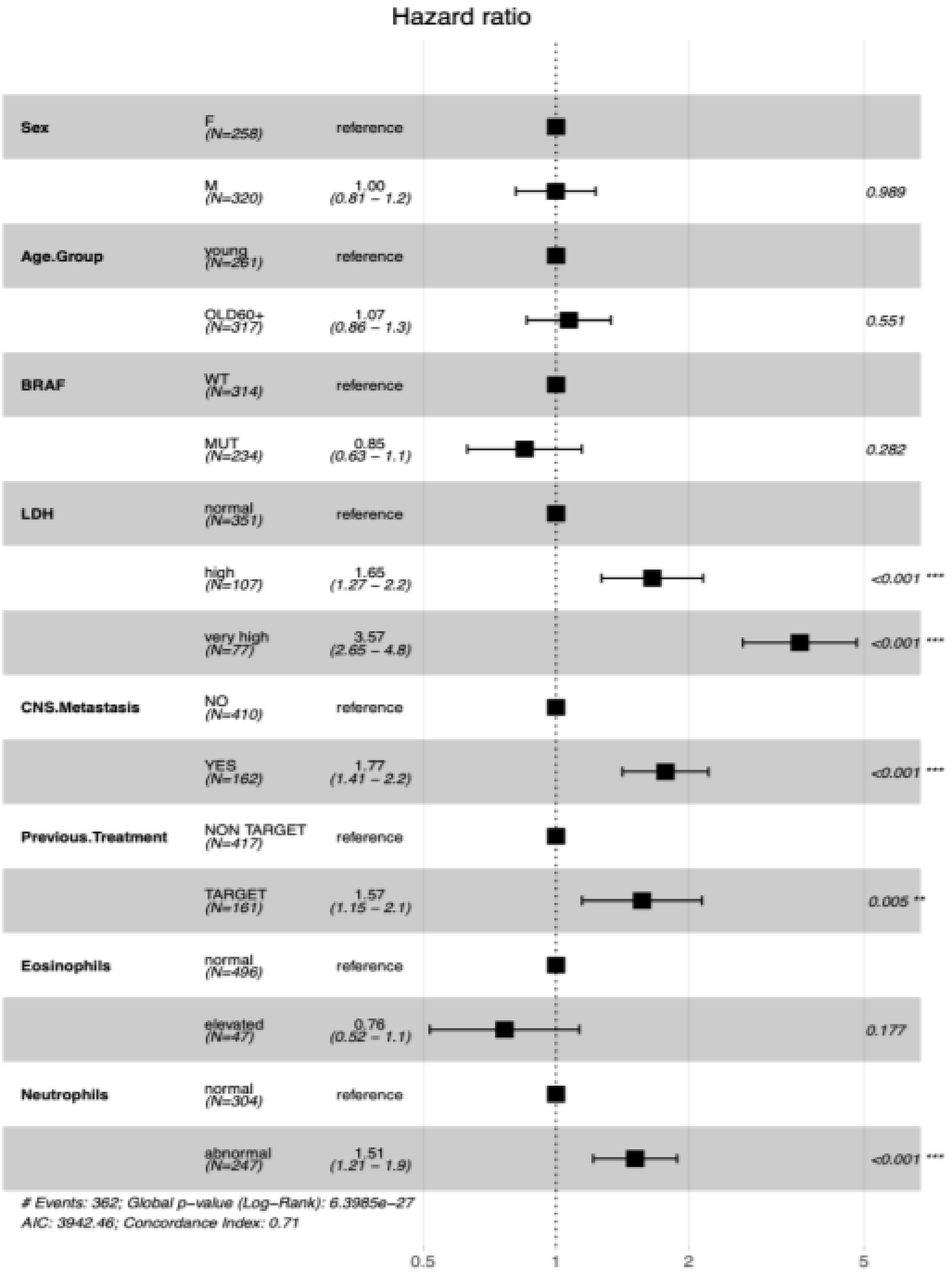


Supplementary Figure 1. Forest plot of multivariate Cox proportional hazards regression results using eight clinical variables from Table 1 as covariates in the model. The first column shows the covariate category name, with the second column indicating the category subgroups and the numbers of patients in each category. The third column shows the hazards ratios of the subgroups when compared to reference category together with their confidence intervals. The confidence intervals are also illustrated by horizontal bars in a logarithmic scale.

#### 4.2 10-fold cross-validation of metastatic melanoma SRF-model

The final SRF model for the 578 patient cohort of metastatic melanoma patients was further validated using 10-fold cross-validation. Folds for cross-validation were generated using the R package caret which produces groups that were balanced by status [12]. Each of the ten cross-validation folds were thus composed of a training set containing 90% of the data and a test set containing the remaining 10%. For each of the ten sets of data, an SRF model trained using its training data and the tuned model parameters from the existing model that was generated using the entire dataset. For each of these 10 models, survival probability predictions at 12, 24, 36 and 60 months were then made using the models test dataset, while risk groups were assigned using the same cut-off values as previously (High-risk: 5-year survival probability <= 0.2; Medium-risk: 5-years survival probability > 0.2 and < 0.41; Low-risk: 5-year survival probability >= 0.41). Average AUC values were calculated across the 10 cross-validation results and an averaged ROC plot was produced using the R package ROCR (Supplementary Figure 2) [13]. Risk group predictions for the 10 cross-validation folds were combined, and a Kaplan-Meier curve was generated with all patients stratified into the three risk categories (Supplementary Figure 3).

#### 4.3 Validation using an external melanoma cohort

Further validation of the SRF-CLICAL model for melanoma was carried out using an external validation cohort composed of 117 metastatic melanoma patients. Values for each variable used in the existing SRF model were extracted from the external-cohort data and variable names were harmonized to match the existing model. The melanoma SRF model was then used to predict survival probabilities of each patient in the external validation cohort at the 12, 24, 36 and 60 month time points. Time-dependent ROC curves at each of the four time points were then generated using the prediction, performance and plot.performance functions from the R package ROCR (Supplementary Figure 4). Each patient in the external validation cohort was then categorized into the three risk groups, using the same survival probability cutoffs as before, and a Kaplan-Meier curve was produced (Supplementary Figure 5).

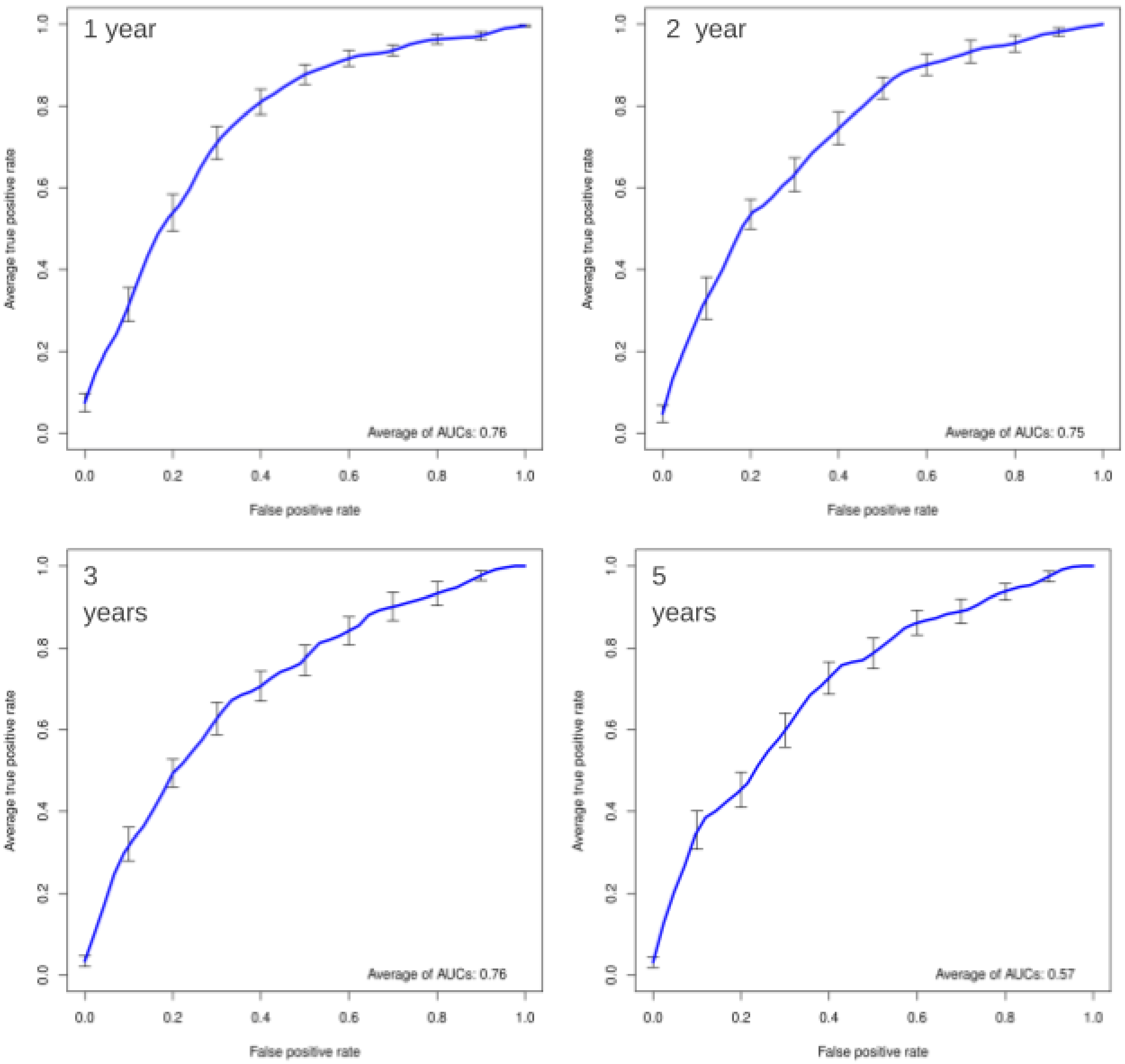


Supplementary Figure 2. Time-dependent receiver operating characteristic (ROC) curves from the 10-fold cross-validation combined results. ROC curves are shown at time points 12, 24, 36 and 60 months. Y-axis in the plots is displaying the True Positive rate (TPR), i.e. Sensitivity, whereas X-axis in the plots is displaying the False Positive Rate (FPR).

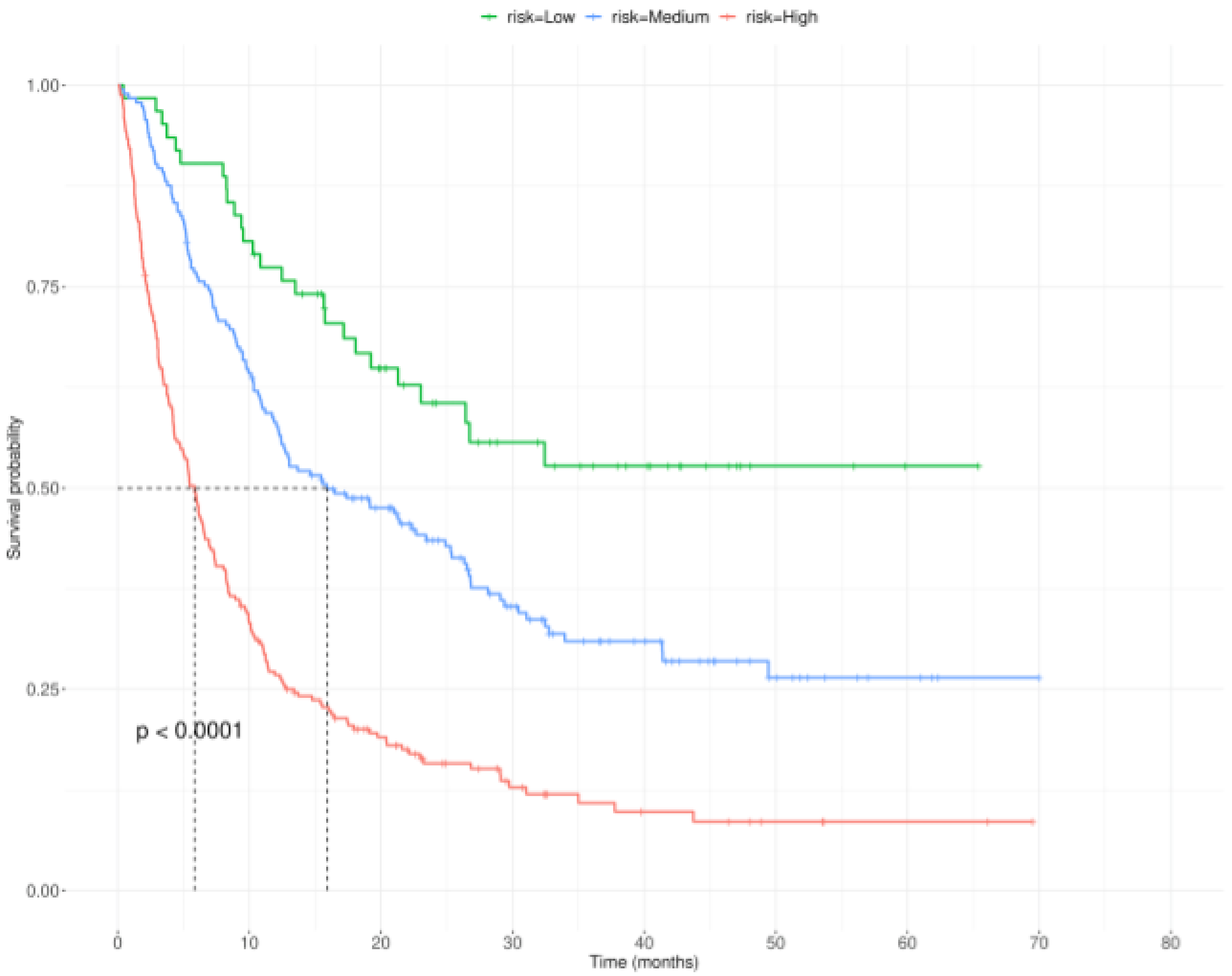


Supplementary Figure 3. Kaplan-Meier survival curves produced using the 10-fold cross-validation survival probability predictions of all 578 melanoma patients divided into three risk groups based on the survival probability predictions. The X-axis shows the survival time in months and Y- axis shows the survival probability estimated based on the observed and censored events.

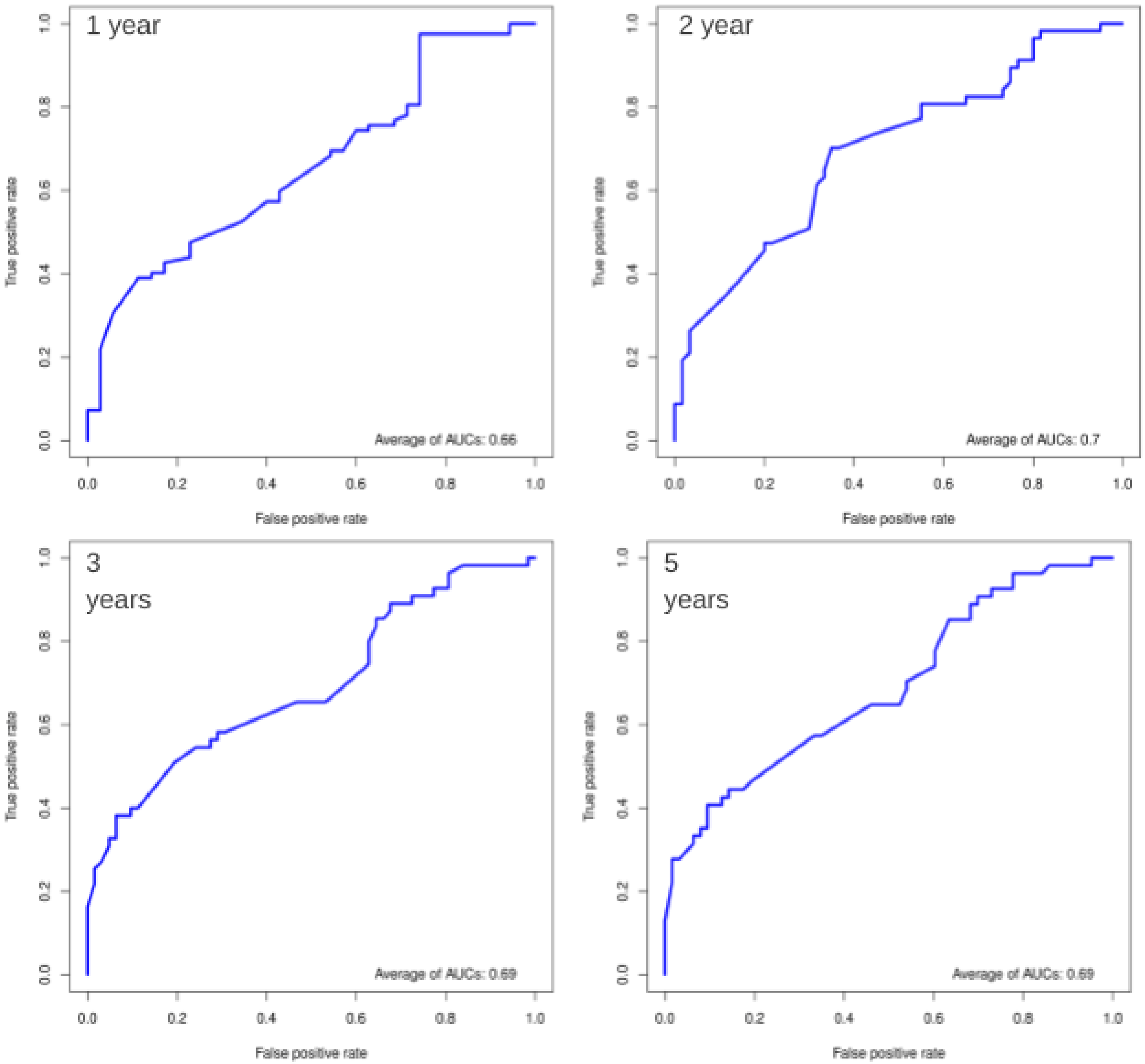


Supplementary Figure 4. Time-dependent receiver operating characteristic (ROC) curves from results of the validation using the external cohort. ROC curves are shown at time points 12, 24, 36 and 60 months. Y-axis in the plots is displaying the True Positive rate (TPR), i.e. Sensitivity, whereas X-axis in the plots is displaying the False Positive Rate (FPR).

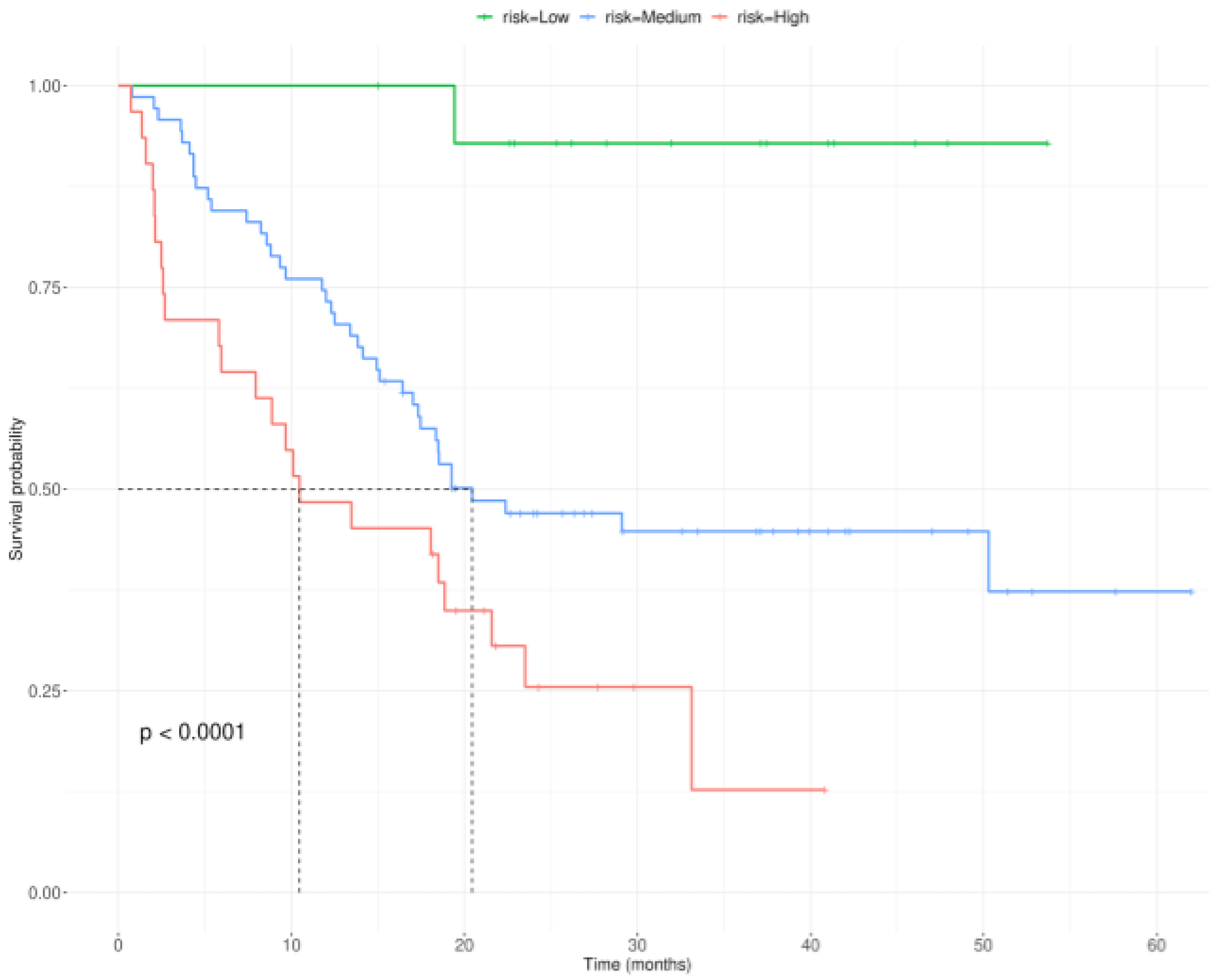


Supplementary Figure 5. Kaplan-Meier survival curves produced using the survival probability predictions from all 117 patients belonging to the external validation cohort. Patients were divided into three risk groups based on the survival probability predictions. The X-axis shows the survival time in months and Y- axis shows the survival probability estimated based on the observed and censored events.

#### 4.4 Analysis of a colorectal cancer cohort using SRF-CLICAL

In order to demonstrate that the SRF-CLICAL approach is robust and easily generalizable, a further proof-of-concept study was conducted using a colorectal cancer cohort comprising 520 patients. Altogether, 14 clinical variables were available for patients in this cohort (Supplementary Table 3). Univariate Cox proportional hazards (Cox PH) regression models were fitted for all 14 variables listed in Supplementary Table 3 with survival time being used as the outcome variable. Effron approximation was used for handling tied death times. The p-values and hazard ratios of the models were inspected to compare the predictive abilities of the independent variables. A survival random forest (SRF) model was then computed for the data using the following ten variables as features: Gender, Age, HLA-A2 genotype, MSI, HER-3, CD8+ IM, HLA-G phenotype, Duke's clinical stage, Histologic differentiation degree and Treatment. Training and optimisation of the SRF model was performed using the same approach described in section 4.1. Using the optimised SRF model, survival probabilities of each patient at 24, 36 and 60 months were predicted with predictSurvProb function of the pec R package. Distribution of the predicted survival probabilities at 5 years (60 months) were examined and used for defining the following risk categories: patients with survival probability < 0.55 were categorised as High risk, patients with survival probability => 0.75 were categorised as Low risk and patients with survival probability in between these thresholds were categorised as Medium risk. Time-dependent receiver operating characteristic (ROC) curves at time points 24, 36 and 60 months were then generated (Supplementary Figure 6). The AUC values at all four time points were quite similar to each other, roughly 70. Lastly, a Kaplan-Meier curve was generated with all patients stratified into the three risk categories (Supplementary Figure 7). The survival curves for all three risk categories showed quite distinct patterns and reached statistical significance.

Furthermore, we carried out 10-fold cross validation of the final optimised SRF-model for colorectal cancer risk prediction. As carried out for the metastatic melanoma cohort, the R package caret was used to produce 10 validation groups that were balanced by status, with each of these ten cross-validation folds being thus composed of a training set containing 90% of the data and a test set containing the remaining 10%. An SRF-model was trained using the training data from each of the ten cross-validation folds, and using the final optimised SRF-model parameters. For each of these 10 newly generated models, survival probability predictions at 12, 24, 36 and 60 months were made using the test set. Risk groups were then assigned using the same cut-off values as previously (High-risk: 5-year survival probability <= 0.55; Medium-risk: 5-years survival probability > 0.55 and < 0.75; Low-risk: 5-year survival probability >= 0.75). Average AUC values were calculated across the 10 cross-validation results and an averaged ROC plot was produced using the R package ROCR (Supplementary Figure 8). Risk group predictions for the 10 cross-validation folds were combined, and a Kaplan-Meier curve was generated with all patients stratified into the three risk categories (Supplementary Figure 9). The cross-validation results were consistent with the results of the original colorectal cancer SRF-model.

Supplementary Table 3. Clinical variables available for the 520 patient colorectal cancer cohort.

| Variable no | Variable name | Groups |
|---|---|---|
| 1 | Age | Old (> 65), Young (<=65) |
| 2 | Gender | Male, Female |
| 3 | Duke’s clinical stage | B, C |
| 4 | Histologic differentiation degree | low, medium, high |
| 5 | HLA-A2 genotype | 0 (no), 1 (yes) |
| 6 | HLA-G genotype | negative (+luminal), positive |
| 7 | Mismatch repair (MMR) deficiency | ProfMMR (proficient), DefMMR (deficient) |
| 8 | HER-3 | low (1), high (2) |
| 9 | CD8+ immunoscore | CD8SCORE1, CD8SCORE2, CD8SCORE3, CD8SCORE4 |
| 10 | CD8+ IT (infiltration in tumour) | 0, 1, 2, 3 |
| 11 | CD8+ IS (infiltration in stroma) | 0, 1, 2, 3 |
| 12 | CD8+ IM (infiltration in margin) | 0, 1, 2, 3 |
| 13 | MHC | low (0, 1, 2), high (3, 4) |
| 14 | Tumor location | DX, SIN |

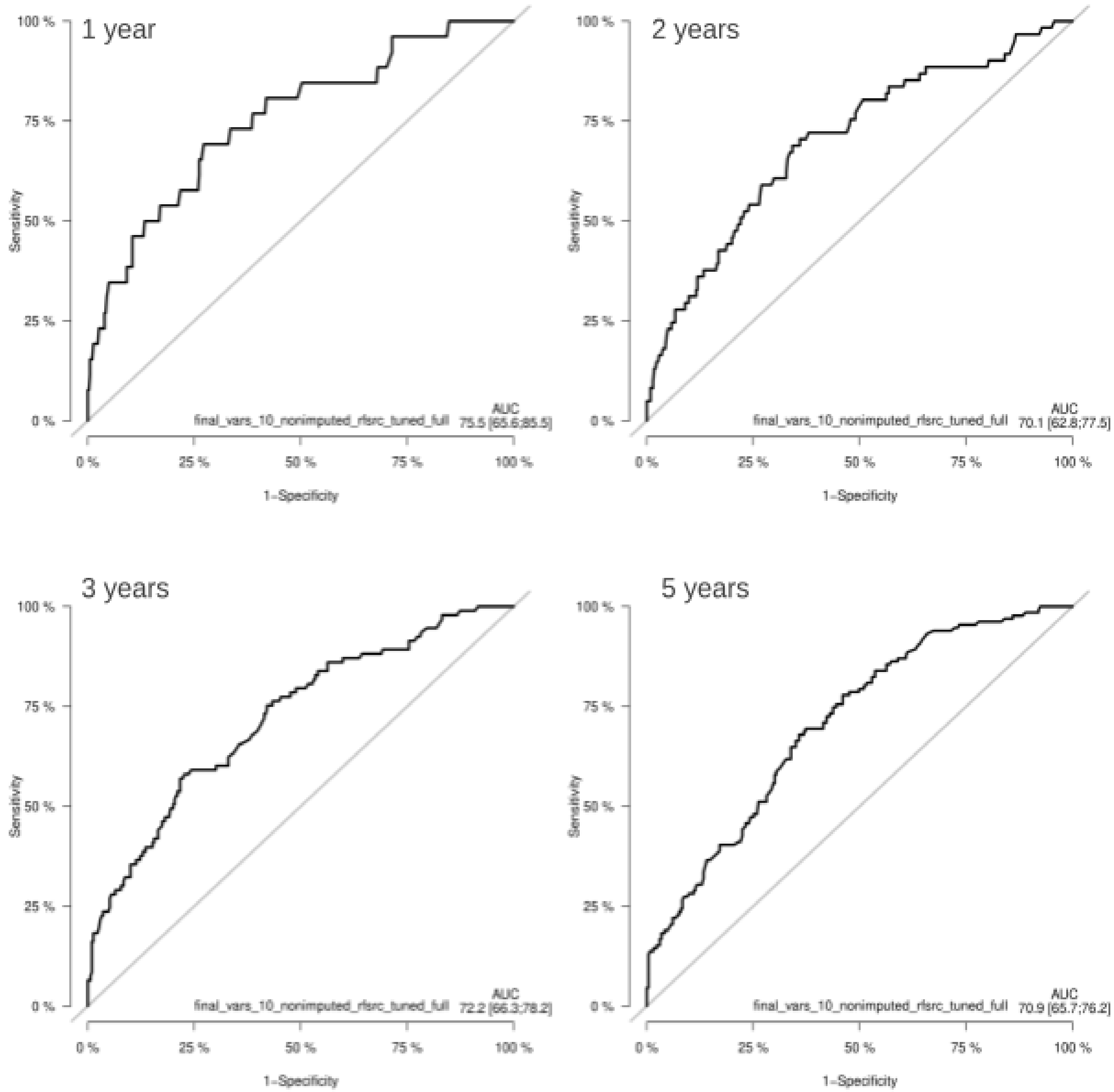


Supplementary Figure 6. Time-dependent receiver operating characteristic (ROC) curves from Colorectal Cancer SRF-model. ROC curves are shown at time points 12, 24, 36 and 60 months. Y-axis in the plots is displaying the True Positive rate (TPR), i.e. Sensitivity, whereas X-axis in the plots is displaying the False Positive Rate (FPR).

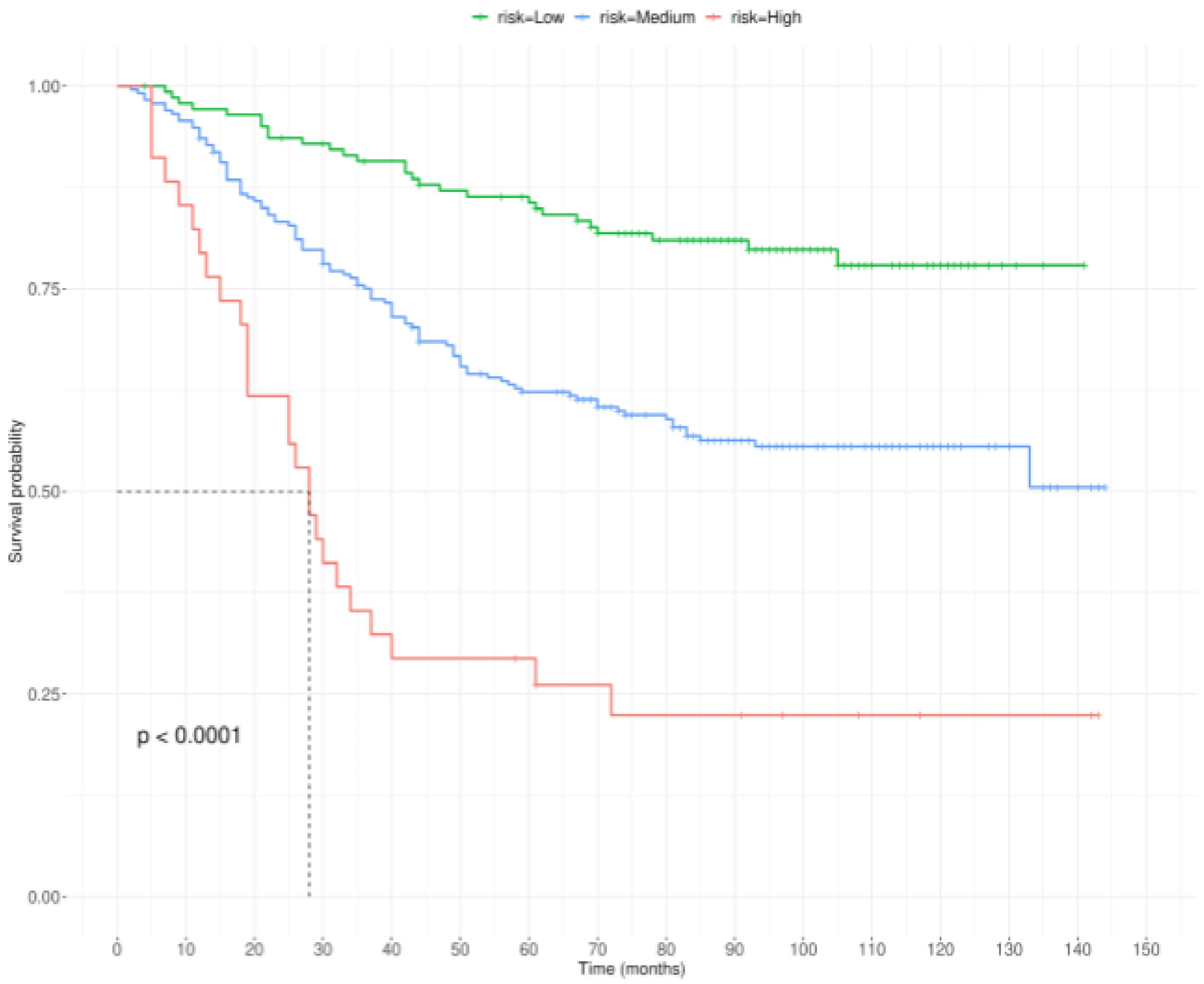


Supplementary Figure 7. Kaplan-Meier survival curves of the colorectal cancer patients divided into three risk groups based on the survival probability predictions obtained using the optimised survival random forest model. Patients were divided into three risk groups based on the survival probability predictions. The X-axis shows the survival time in months and Y- axis shows the survival probability estimated based on the observed and censored events.

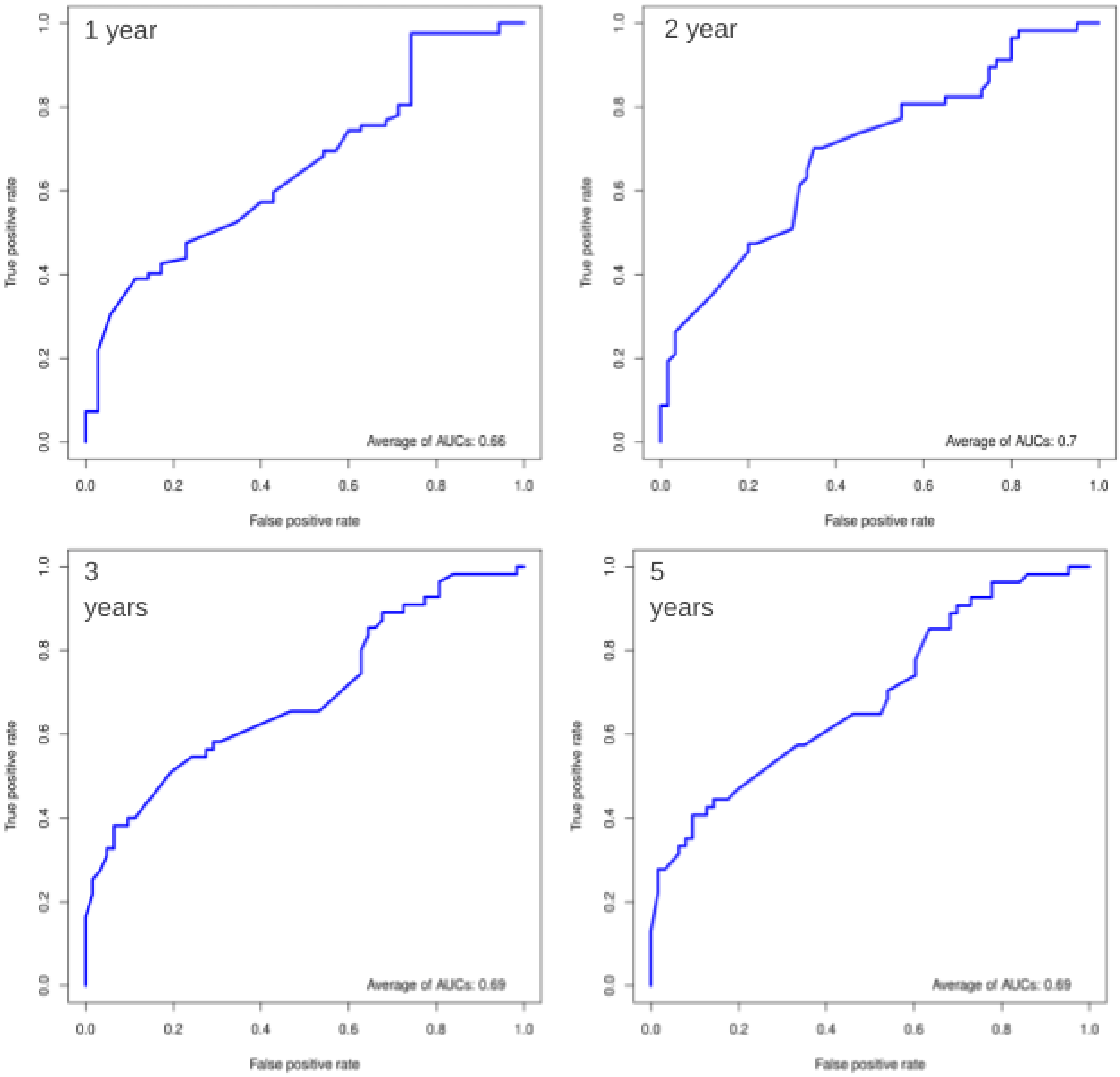


Supplementary Figure 8. Time-dependent receiver operating characteristic (ROC) curves from the 10-fold cross-validation combined results of the colorectal cancer cohort. ROC curves are shown at time points 12, 24, 36 and 60 months. Y-axis in the plots is displaying the True Positive rate (TPR), i.e. Sensitivity, whereas X-axis in the plots is displaying the False Positive Rate (FPR).

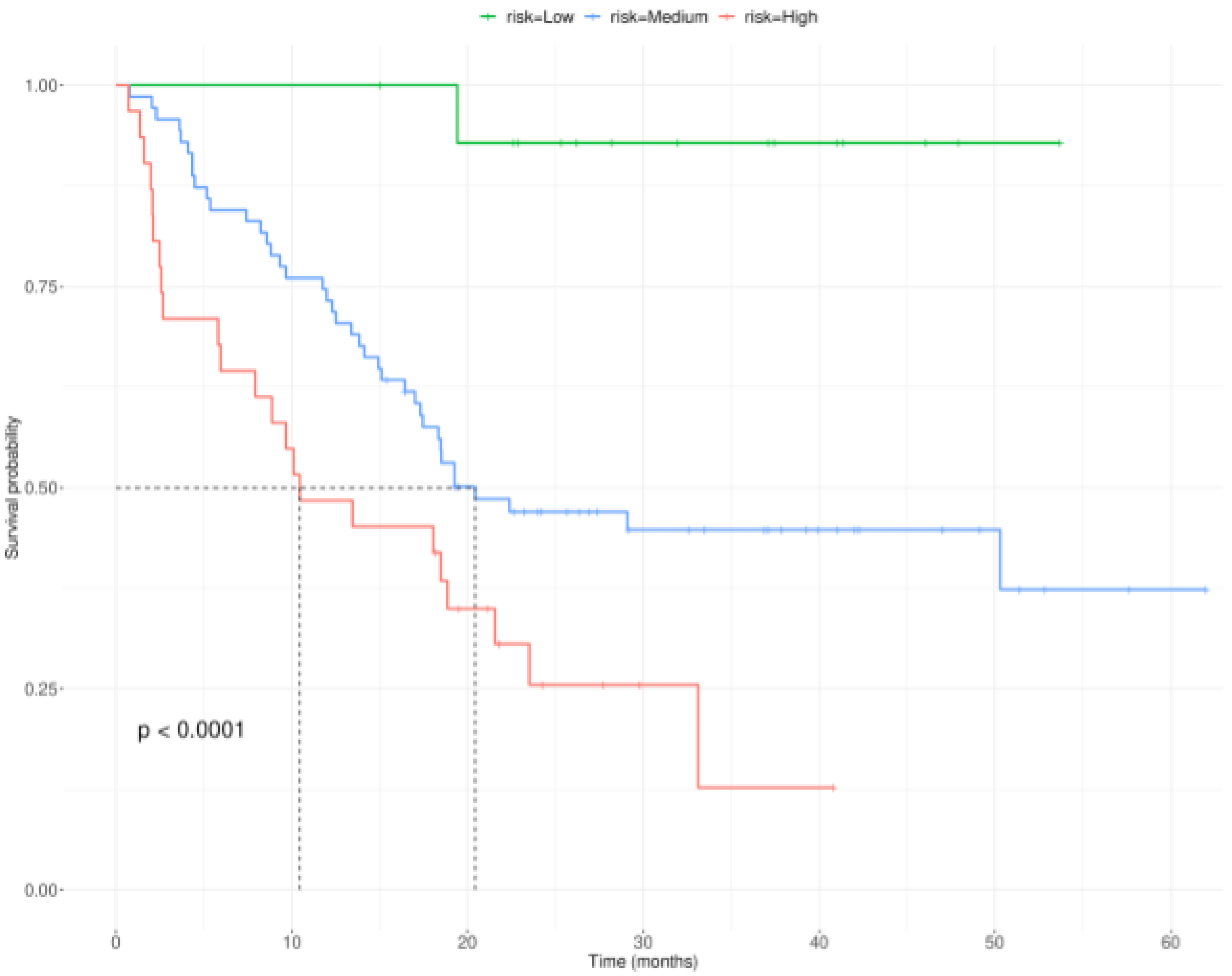


Supplementary Figure 9. Kaplan-Meier survival curves produced using the 10-fold cross-validation survival probability predictions of all colorectal cancer patients divided into three risk groups based on the survival probability predictions. The X-axis shows the survival time in months and Y- axis shows the survival probability estimated based on the observed and censored events.
